# Efferent retargeting in above-knee amputees is positively related to phantom limb pain

**DOI:** 10.1101/2020.08.12.248518

**Authors:** Amanda S. Therrien, Cortney Howard, Laurel J. Buxbaum

**Author notes:** **Please address correspondence to:** Amanda S. Therrien, PhD Moss Rehabilitation Research Institute 50 Township Line Rd. Elkins Park, PA, 19027.

## Abstract

Many individuals who undergo limb amputation experience persistent phantom limb pain (PLP). The underlying mechanism of PLP is unknown, but the phenomenon has been associated with reorganization in sensorimotor cortex following amputation. The traditional view is that cortical reorganization degrades the missing limb’s representation. However, recent work suggests that an amputated limb’s cortical representation remains intact and that reorganization reflects a retargeting of efferent projections to residual muscles proximal to the amputation site. Evidence of retargeting has only been shown in individuals with upper limb amputations, and the relationship of retargeting to PLP is controversial. This study assessed retargeting and its relationship to PLP in 10 individuals with lower limb amputations. We recorded electromyographic (EMG) activity in a residual thigh muscle (vastus lateralis, VL) in patients with above-knee amputations during cyclical movements of the foot. VL activity on the amputated side was compared to that recorded on patients’ intact side while they moved their phantom and intact feet, respectively. VL activity in the patient group was also compared to VL activity from a sample of 9 control participants with no amputation. We show that phantom foot movement is associated with greater VL activity in the amputated leg than that seen in the intact leg as well as that exhibited by controls. The magnitude of residual VL activity was also positively related to ratings of PLP. These results provide the first support for retargeting in lower limb amputees and suggest that retargeting is related to the experience of phantom pain.

**New and Noteworthy:** Previous work has only examined retargeting in upper limb amputees. This study provides evidence for retargeting in lower limb amputees and suggests that retargeting is related to phantom limb pain.

## Introduction

Up to 90% of amputees report experiencing pain in their missing limb following amputation (Kooijman et al. 2000, Jensen et al. 1983). This phantom pain can range from short bouts of moderate pain to chronic, debilitating pain in the missing limb (Hill 1999). Several factors are known to influence the occurrence and extent of phantom pain, such as pre-amputation pain, pre-amputation limb dominance, amputation level, gender, and prosthetic use (Kooijman et al. 2000); however, the precise neural mechanism underlying phantom limb pain remains unknown.

One hypothesized mechanism of phantom limb pain stems from observations of referred sensation of the missing limb to other body segments. For example, stimulating of regions of the face has been shown to elicit phantom sensations in an amputated hand (Ramachandran and Hirstein 1998). Referred sensation may result from neuroplastic changes in sensorimotor cortex following amputation, where cortical regions representing the missing limb begin responding to stimulation of body parts occupying adjacent regions of the sensorimotor somatotopy. Cortical reorganization after amputation has been shown in both humans and animal models in primary somatosensory (Merzenich et al. 1984, Flor et al. 1998, Jain et al. 1998) and motor cortices (Donoghue & Sanes 1988, Cohen et al. 1991, Pasqual-Leone et al. 1996, Wu & Kaas 1999, Qi et al. 2000). The degree of cortical reorganization after amputation has also been shown to positively correlate with ratings of phantom limb pain (Flor et al. 1995, Karl et al. 2001, Lotze et al. 2001).

The prevailing view is that post-amputation cortical changes occur at the expense of the missing limb’s representation. However, there is growing evidence that the cortical representation of an amputated limb remains intact. When moving their phantom hand, upper limb amputees activate regions of sensory and motor cortex homologous to those activated during movements of the intact limb (Makin et al. 2013, Kikkert et al. 2018, Lotze et al. 2001, Roux et al. 2001, 2003). Furthermore, transcranial magnetic stimulation (TMS) of the motor cortex contralateral to the amputated arm can elicit sensations of movement in the phantom hand (Hess et al. 1986, Cohen et al. 1991, Pasqual-Leone et al. 1996), including movements that amputees are unable to generate voluntarily (Mercier et al. 2006).

How can cortical reorganization coexist with a maintained representation of an amputated limb? One prominent hypothesis posits that efferent motor projections remain intact after amputation but are retargeted to muscles proximal to the amputation site (Reilly and Sirigu 2008). Although patients experience sensation in a missing limb, phantom movements can elicit distinct patterns of electromyographic (EMG) activity in residual muscles proximal to the amputation site (Reilly et al. 2006, Gagné et al. 2009). Notably, the residual muscles activated by phantom movement did not contribute substantially to the tested actions prior to amputation (Gagné et al. 2009, Reilly et al. 2006). In patients with above-elbow amputations, residual muscle EMG can be elicited by TMS of the missing hand region suggesting overlap in corticomotor representations (Mercier et al. 2006).

Efferent retargeting after amputation may serve to close the feedback loop necessary to maintain a sensorimotor representation of the missing limb (Reilly and Sirigu 2008) and there is evidence to suggest that a maintained representation may mitigate phantom pain sensations. Better sense of control over phantom limbs is associated with lower ratings of phantom limb pain (Kikkert et al. 2017). Additionally, enhanced control over residual muscles during operation of myoelectric prostheses has been found to decrease phantom pain ratings (Hunter et al. 2008, Lotze et al. 1999, Weiss et al. 1999). Contrary to this idea however, Gagné et al. (2009) found that the degree of EMG modulation in residual arm muscles during movement of a phantom hand positively correlated with ratings of phantom pain. This result highlights the need for further research studying the relationship between retargeting and phantom limb pain.

Previous evidence of retargeting after amputation has only been shown in upper limb amputees. Here, we examined EMG activity in the vastus lateralis (VL) – a knee-extensor muscle in the thigh – in patients with above-knee amputations while they made cyclical movements with their foot. We compared VL activity on the amputated side to that in patients’ intact leg while they moved their phantom and intact feet, respectively. We also compared VL activity in the patient group to recorded that from a sample of control participants with no amputation. We show that phantom foot movement elicits greater residual VL activity than that seen in the intact leg as well as greater activity than that exhibited by controls. Additionally, we show that the magnitude of residual VL activity is positively related to ratings of phantom limb pain. These results provide the first support for efferent retargeting in lower limb amputees and suggest that this process is related to the experience of phantom pain.

## Method

### Participants

10 patients with above-knee amputations and intact vastus lateralis muscle bellies who reported experiencing Phantom Limb Pain (PLP) were recruited from Moss Rehab in Elkins Park, Pennsylvania (4 females, 6 males, mean age ± SD: 56.3 ± 10.8 years, mean education level ± SD: 14.2 ± 3.6 years). In addition 9 control participants, matched to PLP patients for age and education level were recruited from the Neuro-Cognitive Rehabilitation Research Registry at the Moss Rehabilitation Research Institute (4 females, 5 males, mean age ± SD: 57.9 ± 11.0 years, mean education level ± SD: 16.1 ± 3.1 years; Schwartz et al. 2005). More detailed demographic information about the PLP and control participants are shown in Table 1. No participants had a history of psychosis, neurologic disorder, traumatic brain injury or drug abuse. In compliance with the guidelines of the Institutional Review Board of Einstein Healthcare Network, all participants gave informed consent and were compensated for travel expenses and participation.

**Table 1.**
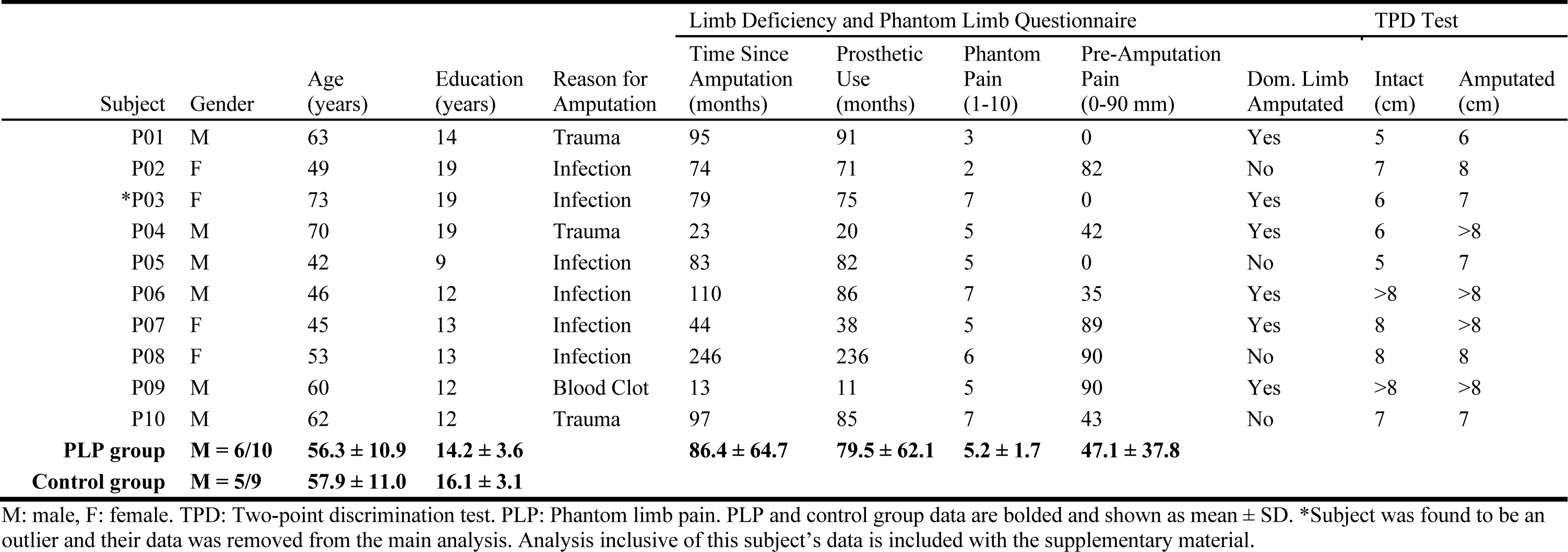
Subject demographics

### Procedure

#### Limb Deficiency and Phantom Limb Questionnaire

The PLP group completed an abbreviated version of the Limb Deficiency and Phantom Limb Questionnaire (Goller et al. 2013). This brief questionnaire assessed prosthesis usage, pre-amputation limb dominance and non-painful phantom limb sensations, including perceived phantom limb size, ability to move the phantom, and level of pain in the limb prior to amputation. Average pain prior to amputation was assessed using a 90mm horizontal line scale that patients were asked to mark with a vertical line: furthest to the left (i.e. 0) indicating no pain, furthest to the right (i.e. 90) indicating the worst pain imaginable. The mark was then measured in mm from the left start of the line. PLP patients also rated their average phantom limb pain using a scale running from 1-10, where 1 indicated very mild pain and 10 indicated completely debilitating pain.

#### Two-Point Discrimination Test

To assess tactile sensation in the thighs on the intact and amputated side, PLP patients completed a modified version of the two-point discrimination test (TPD) included in the Rivermead Assessment of Somatosensory Performance (RASP) (Halligan et al. 1997). The neuro-discriminator used in the RASP is not large enough to yield distances between points that are distinguishable on the thigh so we modified the TPD test to use a caliper and tested the more sensitive inner portion of the thigh. Subjects were asked to close their eyes and lay in a supine position. The initial distance on the caliper was set to 8 cm. On each trial, either one or both points of the caliper were applied to the skin surface and depressed approximately 5 mm. The calipers were held briefly in position before being removed. Following caliper removal subjects were asked to indicate whether they felt that one or two points had been applied to their thigh. A total of 8 trials were performed for each caliper distance, comprising 6 trials with two points applied and 2 trials with only one point applied. The order of these trials was randomized. If the subject’s discrimination over the 8 trials achieved a minimum accuracy of 75% (i.e. 6/8 trials correct), the caliper distance was reduced by 1 cm. The test was then repeated until a caliper distance was reached where the subject could not distinguish the stimuli with at least 75% accuracy. The smallest distance that subjects could distinguish accurately was recorded. If a subject could not reliably distinguish the initial 8 cm distance, a value of “> 8 cm” was recorded.

#### Experimental Task

EMG sensors were placed on the skin of the thigh over the VL muscle in a bipolar configuration, parallel to the direction of the underlying muscle fibers. EMG data was collected using the Trigno™ Wireless System by Delsys (Natick, MA). Sensor locations were shaved if necessary.

EMG activity was recorded from the VL while subjects performed cyclical movements with the ipsilateral foot. Subjects performed the foot movements while lying supine with their eyes closed. EMG activity was recorded for 2 movement conditions: ankle flexion/extension and ankle abduction/adduction. A single trial involved performing the cyclical movement for 20 s. Subjects performed 3 trials for each movement condition and each leg for a total of 12 trials per subject. Prior to beginning the experimental task, EMG activity during a maximum voluntary contraction (MVC) of the VL on each leg was recorded.

### Analysis

Preliminary data analysis revealed that one subject in the PLP group was an extreme outlier (P03, the normalized EMG amplitude on the amputated leg was >2.5 SD of the PLP group mean for both movement conditions). Their data were excluded from the analyses reported here. However, to ensure that excluding this participant did not bias our results, analyses including this participant’s data are shown in the supplementary material.

Measures from the modified Limb Deficiency and Phantom Limb Questionnaire and TPD test were submitted to Spearman’s rank order correlations to assess their relationship with amputees’ reported average phantom limb pain. For TPD measures of > 8cm, a value of 9 cm was entered into the correlation. Correlation results informed what measures were necessary to incorporate into mixed-effect linear regression models that assessed the relationship between EMG activity and phantom limb pain. Because correlations were not interpreted independently, no correction for multiple comparisons was performed.

All EMG data were rectified and band-pass filtered at 20-2000 Hz. The time series of EMG amplitudes from each movement trial were then normalized as ratios of the mean EMG amplitude obtained during maximum voluntary contraction (MVC) of the VL on each side. The mean amplitude from the normalized time series was then obtained for each trial.

Statistical analysis comprised a series of mixed-effects linear regression models using LMER4 (Bates et al. 2015) performed with subject means of normalized EMG amplitude as the dependent variable. To control for individual differences in EMG amplitude, participant-specific intercepts were entered as random predictors. The categorical variable of pre-amputation limb dominance (coded as yes or no), which could not be included in the correlation analysis described above, was entered as a fixed-effect in all models.

The first regression model tested for VL recruitment during movement of the foot. PLP patients’ limb status was coded as “amputated” or “intact.” Limb status in the control group was pseudo-randomly assigned as “amputated” or “intact” with the constraint that 5 of 9 controls had their dominant limb assigned as amputated (to match the PLP group, see Table 1). Limb status (amputated, intact), group (PLP, control), and an interaction term of group and limb were entered into the model as predictors of interest. Controlling variables included the fixed effect predictor of pre-amputation limb dominance (yes, no), and the random intercept of participant.

The second regression model tested the relationship between scaled EMG amplitude and phantom limb pain within the PLP group only. Limb status (amputated, intact), average phantom limb pain (1-10), and an interaction term of average phantom limb pain and limb status were entered into the model as predictors of interest. Controlling variables included the fixed effect predictor of dominant limb amputation (yes, no), and the random intercept of participant.

Statistical significance of fixed-effect predictors was assessed using likelihood ratio tests. To analyze interactions, the data were split along each predictor included in the interaction, and likelihood ratio tests were performed to assess the significance of the other predictor. An alpha value of 0.05 was used for all analyses.

## Results

The PLP group’s responses to the Limb Deficiency and Phantom Limb Questionnaire and results of the TPD testing are included in Table 1. No significant relationships were found between any of these measures and average phantom limb pain rating (Table 2). As a result, none of these measures were included as predictors in either regression model.

**Table 2.** Relationship among measures of limb deficiency and TPD and average phantom pain rating.

|  | Months post<br>amputation | Prosthetic<br>use | Pre-amputation<br>pain | TPD intact<br>limb | TPD amputated<br>limb |
| --- | --- | --- | --- | --- | --- |
| Spearman's Rho (*r*) | .542 | .420 | .106 | .467 | .194 |
| Sig. (*p*) | .132 | .261 | .786 | .205 | .618 |

Correlations were analyzed using Spearman's Rank Order Correlations with an alpha of 0.05.

Regression analyses showed that movement condition was a significant fixed-effect predictor of EMG amplitude. That is, the flexion/extension condition was associated with greater EMG amplitude compared with the abduction/adduction condition (*χ*2(1) = 11.18, *p* <.001). Given this main effect, further regression analyses were conducted separately for the two movement conditions.

Within the flexion/extension movement condition, significant fixed-effect predictors of EMG amplitude were limb dominance (*χ*2(1) = 4.114, *p*=.043) and limb status (*χ*2(1) = 12.884, *p* <.001). Both limb dominance and performing foot movement on the amputated side were associated with greater EMG amplitudes overall. There was no fixed effect of group (*χ*2(1) = 2.486, *p*=.115); however, there was a significant interaction among group and limb status (*χ*2(1) = 20.389, *p* <.001; Figure 1). To analyze the interaction, the data were first divided by group. Within the PLP group, the amputated leg showed greater EMG amplitudes than the intact leg (*χ*2(1) = 22.253, *p* <.001). Within the control group, there was no effect of limb status (*χ*2(1) = .310, *p*= .578). Dividing the data by limb status revealed that within the intact limb, there was no difference between groups (*χ*2(1) = .212, *p*= .645). However, the PLP group showed greater EMG amplitudes than the control group within the amputated leg (or the leg coded as amputated in the control group; *χ*2(1) = 8.482, *p*= .004).

**Figure 1.**
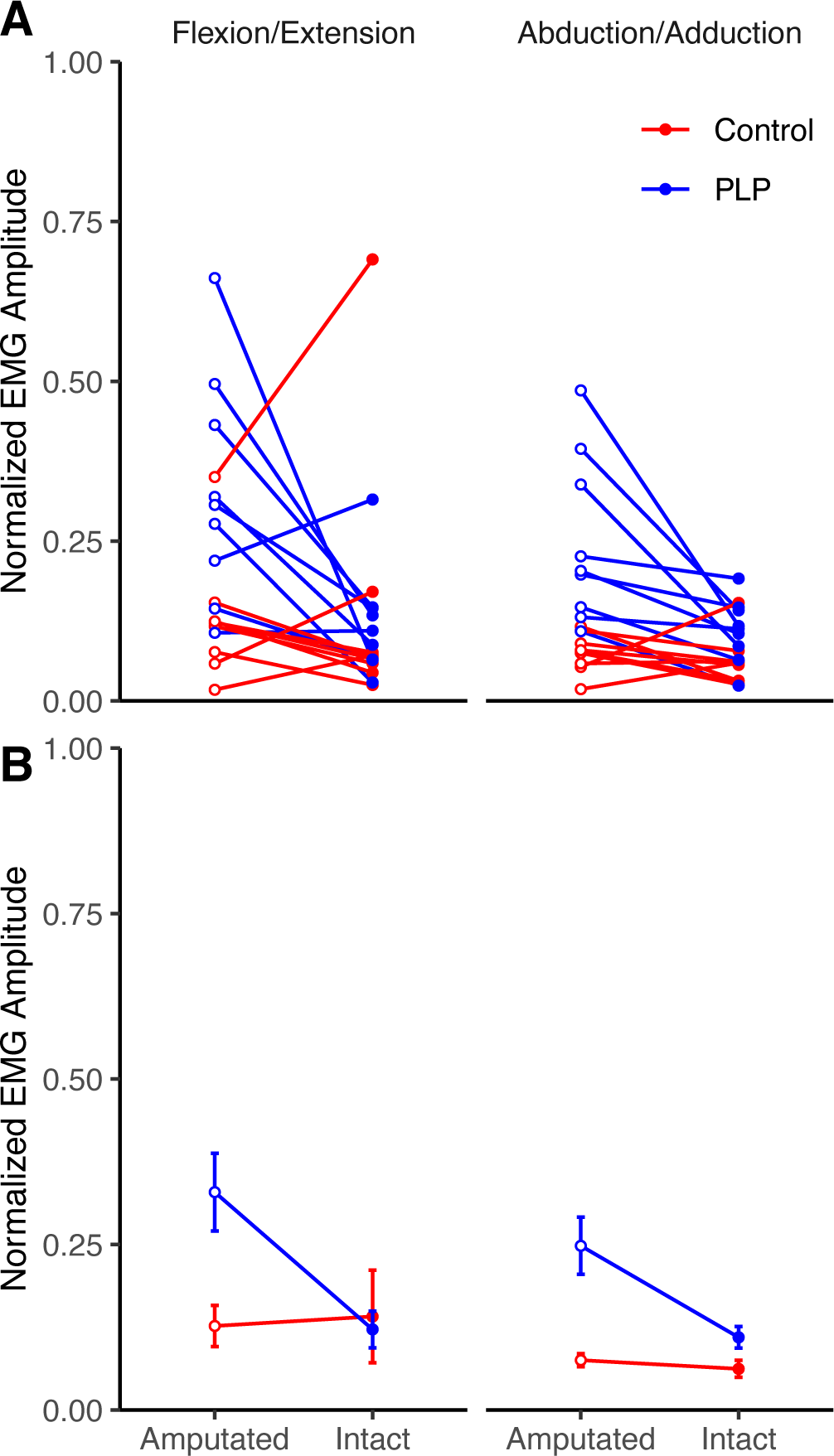
Normalized EMG amplitude during cyclical foot movements. (A) Individual subject data for the two movement conditions. (B) Group mean data showing the interaction among group and limb status for the two movement conditions. PLP patients showed greater EMG amplitudes in their amputated leg compared to their intact leg. There was no effect of limb status in the control group.

Within the abduction/adduction movement condition, significant fixed effect predictors of EMG amplitude were group (*χ*2(1) = 14.099, *p*< .001) and limb status (*χ*2(1) = 18.392, *p*< .001). EMG amplitudes were greater overall in the PLP group compared to controls and in the amputated compared to intact leg. There was no effect of limb dominance (*χ*2(1) = 3.527 *p*= .060). The interaction among group and limb status was significant (*χ*2(1) = 15.164, *p*< .001; Figure 1). Analyzing the interaction showed that within the PLP group, EMG amplitudes were greater in the amputated leg than in the intact leg (*χ*2(1) = 18.245, *p*< .001). There was no effect of limb status within the control group (*χ*2(1) = 1.701, *p*=.192). Analyzing the interaction within limb status showed that group was a significant predictor of EMG amplitude in both the intact (*χ*2(1) = 4.909, *p*= .027) and amputated legs (*χ*2(1) = 12.568, *p*<.001). EMG amplitudes were greater in the PLP group compared to controls.

We used a second regression model to examine the relationship between EMG amplitude and phantom limb pain within the PLP group. In the flexion/extension movement condition significant fixed effect predictors of EMG amplitude were limb dominance (*χ*2(1) = 4.019, *p* = .045), limb status (*χ*2(1) = 24.035, *p* < .001), and average phantom limb pain (*χ*2(1) = 7.784, *p* < .001). Limb dominance was associated with greater EMG amplitude as was performing foot movement in the amputated leg. Greater average phantom limb pain rating predicted greater EMG amplitude overall. Notably, there was a significant interaction between limb status and phantom limb pain rating (*χ*2(1) = 6.525, *p* < .011; Figure 2). Analyzing the interaction showed that greater phantom limb pain was associated with greater EMG amplitudes in the amputated leg (*χ*2(1) = 6.392, *p* = .011), but showed no relationship with EMG amplitude in the intact leg (*χ*2(1) = .238, *p* = .626).

**Figure 2.**
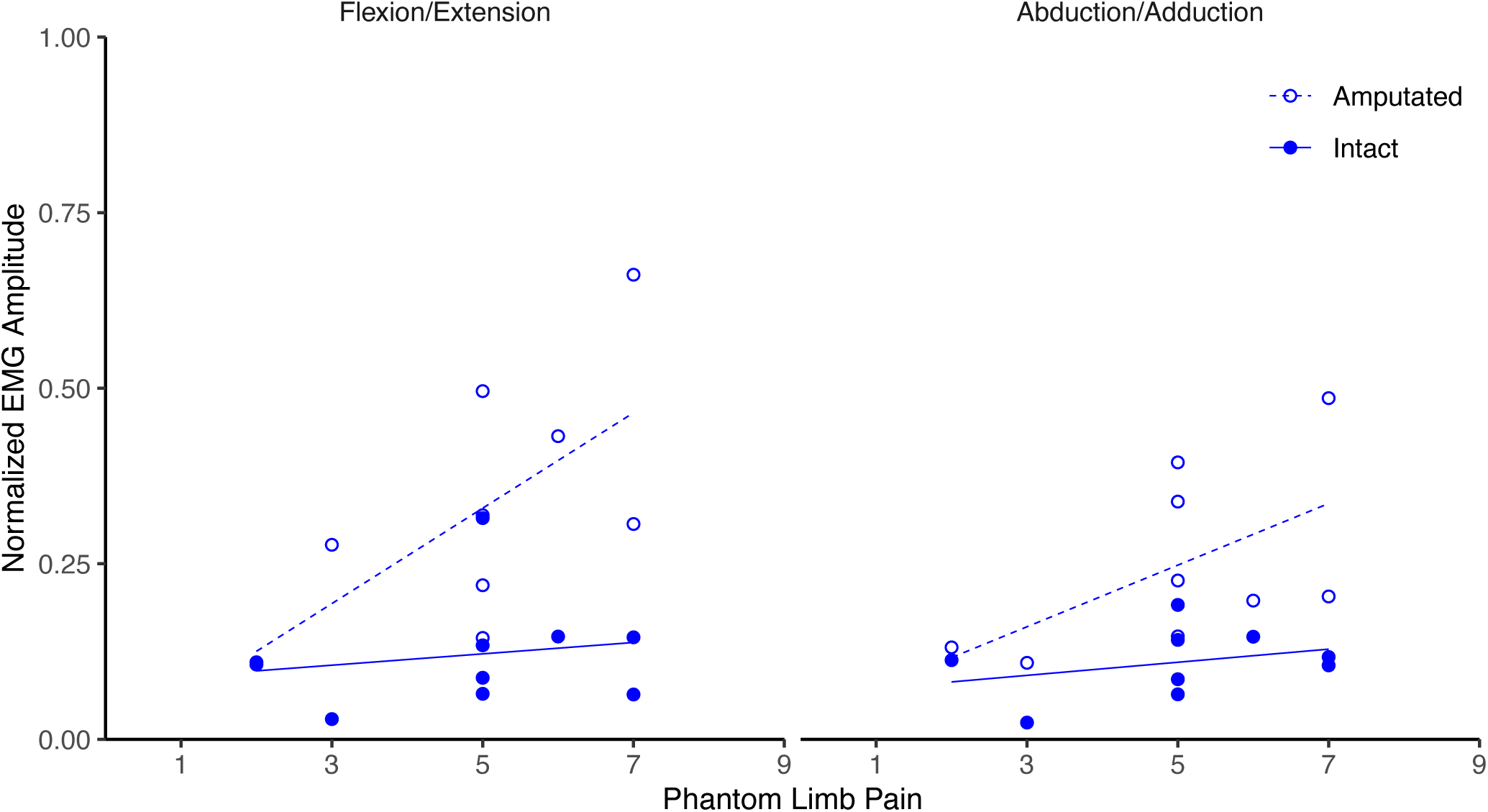
Relationship between normalized EMG amplitude and ratings of phantom limb pain within the PLP group for the flexion/extension (A) and abduction/adduction (B) movement conditions. Plots show two data points per participant corresponding to the amputated and intact legs. Phantom limb pain rating was positively associated with EMG amplitude in the amputated leg, but not in the intact leg.

In the abduction/adduction movement condition, limb dominance, limb status and average phantom limb pain rating were also significant fixed-effect predictors of EMG amplitude. Limb dominance was associated with greater EMG amplitudes ((*χ*2(1) = 4.266, *p* = .039) as was performing the movement on the amputated side (*χ*2(1) = 18.520, *p* < .001). Greater average phantom limb pain rating was associated with greater EMG amplitude overall (*χ*2(1) = 5.277, *p* = .022); however, the interaction between limb status and phantom limb pain rating did not reach significance (*χ*2(1) = 3.570, *p* = .059).

## Discussion

In this study we assessed whether patients with above-knee amputations show evidence of efferent retargeting during movements of their phantom foot and whether efferent retargeting is associated with their experience of phantom limb pain (PLP). We found that cyclical movement of a phantom foot was associated with greater EMG activity in the VL muscle (a knee extensor) of the amputated leg compared to VL EMG in the intact leg during movement of the intact foot. Phantom foot movement also elicited greater VL EMG in the amputated leg compared to that seen in a group of control participants who had no amputation. Within the PLP group, ratings of average phantom foot pain were associated with greater VL EMG and this was specific to the amputated leg when the foot performed flexion/extension movements. Overall, our results provide support for retargeting after amputation in the lower limb and provide evidence of a positive relationship between retargeting and phantom limb pain.

Previous studies of efferent retargeting have only examined persons with upper limb amputations (Reilly et al. 2006, Mercier et al. 2006, Gagné et al. 2009). Our study is the first, to our knowledge, to assess retargeting in individuals with lower limb amputations. Importantly, previous studies have noted key features of retargeting. The first key feature is that phantom movement elicits EMG activity in residual muscles that would not have contributed to movement of the limb prior to amputation. In our study, VL activity in the amputated leg during phantom foot movement was greater than that elicited by foot movement in the intact leg as well as that seen in a group of control participants. This result indicates that phantom foot movement recruited the residual VL to a greater extent than what would have happened normally.

A second key feature of retargeting is that different movements of the phantom limb elicit distinct patterns of EMG in residual muscles. While we did find that movement condition was a significant predictor of VL EMG amplitude, this was the case for both the PLP and control groups. Previous studies noting modulation of EMG based on movement type also recorded EMG from multiple muscles (Reilly et al. 2006; Gagné et al. 2009). The distinct EMG patterns noted in these studies were observed across muscles. As our study only measured EMG in one muscle, we could not attempt to replicate this result.

We found that EMG activity in the residual VL during movement of a phantom foot was positively related to patients’ phantom pain ratings. Gagné et al. (2009) also noted a positive correlation between EMG activity in residual muscles of the upper arm and phantom hand pain. Thus, our results are consistent with this previous work. However, the mechanisms underlying this positive relationship between retargeting and phantom pain remain unclear. One hypothesis concerns incongruence between expected and actual somatosensory feedback during phantom limb movement (Gagné et al. 2009). Retargeting hinges upon the notion that a central representation of the phantom limb remains intact following amputation. As a result, predictive computation of the sensory consequences of phantom limb movement would also be hypothesized to remain intact (Wolpert and Ghahramani 2000). If efferent projections to the phantom limb are altered to target residual muscles not normally involved in movement of that limb, somatosensory signals resulting from phantom limb movement would constantly differ from sensory predictions. Complementary to this notion are observations that phantom limb pain can be mitigated using mirror-box (Chan et al. 2007; Ramachandran and Rogers-Ramachandran 1996) and virtual reality (Ambron et al. 2018) interventions designed to reduce sensory incongruence by providing visual feedback of the missing limb.

Yet, sensory prediction errors are encountered throughout daily life and normally drive an adaptive recalibration of sensorimotor relationships to keep movement and perception attuned to the environment (Krakauer et al. 2000, Shadmehr et al. 2010). Recent work suggests that sensory recalibration may be impaired in persons with amputation (Ortiz-Catalan et al. 2020), which implies a fundamental disruption to the processing of afferent signals in the affected limb even when peripheral afferents survive amputation. A chronic impairment in afferent processing may trigger the development of painful sensations through maladaptive changes to cortical and subcortical regions associated with the pain matrix (Harris 1999). However, further work is needed to discern a precise neural mechanism that could underly such a phenomenon.

An alternative explanation is that a relationship between efferent retargeting and phantom limb pain reflects a spurious correlation resulting from the co-occurrence of two independent processes. Efferent retargeting has been hypothesized to occur through reorganization within primary motor cortex as well as at the peripheral level through regrowth of motor neurons that survive amputation (Reilly and Sirigu 2008). Phantom limb pain has also been posited to result from peripheral mechanisms. Peripheral afferent neurons that survive the amputation undergo sprouting, which can lead to the formation of painful neuromas at the amputation site (Vaso et al. 2014). Phantom limb pain resulting from neuroma formation could occur alongside, but independently of, any efferent retargeting.

In summary we have shown that movement of a phantom foot elicits activity in the residual VL muscle of patients with above-knee amputations that is greater than what is normal for intact foot movement. Additionally, recruitment of the residual VL muscle was positively associated with ratings of phantom foot pain, consistent with previous findings in arm amputations. Overall, our results are consistent with the efferent retargeting hypothesis and are the first to demonstrate this phenomenon in individuals with lower limb amputation. Further work is needed to enhance the characterization of retargeting in this population and determine the precise mechanism underlying the relationship between retargeting and phantom pain.

## Acknowledgements

This work was supported by the Moss Rehabilitation Research Institute.

## Supplementary Materials and Source Data

All supplementary results and source data can be found here: https://github.com/therriea/Retargeting_and_PLP

## Notes

https://github.com/therriea/Retargeting_and_PLP

### Competing Interest Statement

The authors have declared no competing interest.

## References

Ambron E, Miller A, Kuchenbecker K, Buxbaum L, Coslett H. Immersive Low-Cost Virtual Reality Treatment for Phantom Limb Pain: Evidence from Two Cases. Front Neurol. 9: 67, 2018. doi: 10.3389/fneur.2018.00067

Bates D, Mächler M, Bolker B, Walker S. Fitting linear mixed-effects models using lme4. J Stat Soft. 67, 2015. doi: 10.18637/jss.v067.i01

Chan B, Witt R, Charrow A, Magee A, Howard R, Pasquina P, Heilman K, Tsao J. Mirror therapy for phantom limb pain. N Engl J Med. 357: 2206–7, 2007. doi: 10.1056/nejmc071927

Cohen L, Bandinelli S, Findley T, Hallet M. Motor reorganization after upper limb amputation in man: A study with forcal magnetic stimulation. Brain. 114: 615–627, 1991. doi: 10.1093/brain/114.1.615

Donoghue J, Sanes J. Organization of adult motor cortex representation patterns following neonatal forelimb nerve injury in rats. J Neurosci. 8: 3221–3232, 1998. doi: 10.1523/jneurosci.08-09-03221.1988

Flor H, Elbert T, Knecht S, Wienbruch C, Pantev C, Birbaumers N, Larbig W, Taub E. Phantom-limb pain as a perceptual correlate of cortical reorganization following arm amputation. Nature. 375: 482–484, 1995. doi: 10.1038/375482a0

Flor H, Elbert T, Mühlnickel W, Pantev C, Wienbruch C, Taub E. Cortical reorganization and phantom phenomena in congenital and traumatic upper-extremity amputees. Exp Brain Res. 119: 205–212, 1998. doi: 10.1007/s002210050334

Gagné M, Reilly K, Hétu S, Mercier C. Motor control over the phantom limb in above-elbow amputees and its relationship with phantom limb pain. Neuroscience. 162: 78–86, 2009. doi: 10.1016/j.neuroscience.2009.04.061

Goller A, Richards K, Novak S, Ward J. Mirror-touch synaesthesia in the phantom limbs of amputees. Cortex. 49: 243–251, 2013. doi: 10.1016/j.cortex.2011.05.002

Halligan P, Marshall J, Hunt M, Wade D. Somatosensory assessment: can seeing produce feeling? J Neurol. 244: 199–203, 1997. doi: 10.1007/s004150050073

Harris A. Cortical origin of pathological pain. Lancet. 354: 1464–1466, 1999. doi: 10.1016/s0140-6736(99)05003-5

Hess C, Mills K, Murray N. Magnetic stimulation of the human brain: Facilitation of motor responses by voluntary contraction of ipsilateral and contralateral muscles with additional observations on an amputee. Neurosci Lett. 71: 235–240, 1986. doi: 10.1016/0304-3940(86)90565-3

Hill A. Phantom limb pain. J Pain Symptom Manag. 17: 125–142, 1999. doi: 10.1016/s0885-3924(98)00136-5

Hunter J, Katz J, Davis K. Stability of phantom limb phenomena after upper limb amputation: A longitudinal study. Neuroscience. 156: 939–949, 2008. doi: 10.1016/j.neuroscience.2008.07.053

Jain N, Catania K, Kaas J. A histologically visible representation of the fingers and palm in primate area 3b and its immutability following long-term deafferentations. Cereb Cortex. 8: 227–236, 1998. doi: 10.1093/cercor/8.3.227

Jensen T, Krebs B, Nielsen J, Rasmussen P. Phantom limb, phantom pain and stump pain in amputees during the first 6 months following limb amputation. Pain. 17: 243–256, 1983. doi: 10.1016/0304-3959(83)90097-0

Kikkert S, Johansen-Berg H, Tracey I, Makin T. Reaffirming the link between chronic phantom limb pain and maintained missing hand representation. Cortex. 106: 174–184, 2018. doi: 10.1016/j.cortex.2018.05.013

Kikkert S, Mezue M, Slater D, Johansen-Berg H, Tracey I, Makin T. Motor correlates of phantom limb pain. Cortex. 95: 29–36, 2017. doi: 10.1016/j.cortex.2017.07.015

Karl A, Birbaumer N, Lutzenberger W, Cohen L, Flor H. Reorganization of motor and somatosensory cortex in upper extremity amputees with phantom limb pain. J Neurosci. 21: 3609–3618, 2001. doi: 10.1523/jneurosci.21-10-03609.2001

Kooijman C, Dijkstra P, Geertzen J, Elzinga A, Schans C. Phantom pain and phantom sensations in upper limb amputees: an epidemiological study. Pain. 87: 33–41, 2000. doi: 10.1016/s0304-3959(00)00264-5

Krakauer J, Pine Z, Ghilardi M, Ghez C. Learning of visuomotor transformations for vectorial planning of reaching trajectories. J Neurosci. 20: 8916–8924, 2000. doi: 10.1523/jneurosci.20-23-08916.2000

Lotze M, Flor H, Grodd W, Larbig W, Birbaumer N. Phantom movements and pain: An fMRI study in upper limb amputees. Brain. 124: 2268–2277, 2001. doi: 10.1093/brain/124.11.2268

Lotze M, Grodd W, Birbaumer N, Erb M, Huse E, Flor H. Does use of a myoelectric prosthesis prevent cortical reorganization and phantom limb pain? Nat Neurosci. 2: 501–502, 1999. doi: 10.1038/9145

Makin T, Scholz J, Filippini N, Slater D, Tracey I, Johansen-Berg H. Phantom pain is associated with preserved structure and function in the former hand area. Nat Commun. 4: 1570, 2013. doi: 10.1038/ncomms2571

Mercier C, Reilly K, Vargas C, Aballea A, Sirigu A. Mapping phantom movement representations in the motor cortex of amputees. Brain. 129: 2202–2210, 2006. doi: 10.1093/brain/awl180

Merzenich M, Nelson R, Stryker M, Cynader M, Schoppmann A, Zook J. Somatosensory cortical map changes following digit amputation in adult monkeys. J Comp Neurol. 224: 591–605, 1984. doi: 10.1002/cne.902240408

Ortiz-Catalan M, Mastinu E, Bensmaia S. Chronic use of a sensitized bionic hand does not remap the sense of touch. medRxiv, 2020. doi: 10.1101/2020.05.02.20089185

Pascual-Leone A, Peris M, Tormos JM, Pascual AP, Catala MD. Reorganization of human cortical motor output maps following traumatic forearm amputation. NeuroReport. 7: 2068–70, 1996. doi: 10.1097/00001756-199609020-00002

Qi H, Stepniewska I, Kaas J. Reorganization of primary motor cortex in adult macaque monkeys with long-standing amputations. J Neurophysiol. 84: 2133–2147, 2000. doi: 10.1152/jn.2000.84.4.2133

Ramachandran V, Hirstein W. The perception of phantom limbs. The D. O. Hebb lecture. Brain. 121: 1603–1630, 1998. doi: 10.1093/brain/121.9.1603

Ramachandran V, Rogers-Ramachandran D. Synaesthesia in phantom limbs induced with mirrors. Proc Biol Sci. 263: 377–86, 1996. doi: 10.1098/rspb.1996.0058

Reilly K, Sirigu A. The motor cortex and its role in phantom limb phenomena. Neuroscientist. 14: 195–202, 2008. doi: 10.1177/1073858407309466

Reilly K, Mercier C, Schieber, M., Sirigu, A. Persistent hand motor commands in the amputees’ brain. Brain. 129: 2211–2223, 2006. doi: 10.1093/brain/awl154

Roux F, Ibarrola D, Lazorthes Y, Berry I. Virtual movements activate primary sensorimotor areas in amputees: Report of three cases. Neurosurgery. 49: 736–742, 2001. https://dx.doi.org/10.1097/00006123-200109000-00039

Roux F, Lotterie J, Cassol E, Lazorthes Y, Sol J, Berry I. Cortical areas involved in virtual movement of phantom limbs: Comparison with normal subjects. Neurosurgery. 53: 1342–1353, 2003. doi: 10.1227/01.neu.0000093424.71086.8f

Schwartz M, Brecher A, Whyte J, Klein M. A patient registry for cognitive rehabilitation research: A strategy for balancing patients’ privacy rights with researchers’ need for access. Arch Phys Med Rehabil. 86: 1807–1814, 2005. doi: 10.1016/j.apmr.2005.03.009

Shadmehr R, Smith M, Krakauer J. Error correction, sensory prediction, and adaptation in motor control. Neuroscience. 33: 89–108, 2010. doi: 10.1146/annurev-neuro-060909-153135

Vaso A, Adahan H, Gjika A, Zahaj S, Zhurda T, Vyshka G, Devor M. Peripheral nervous system origin of phantom limb pain. Pain. 155: 1384–1391, 2014. doi: 10.1016/j.pain.2014.04.018

Wolpert D, Ghahramani Z. Computational principles of movement neuroscience. Nat Neurosci. 3: 1212–1217, 2000. doi: 10.1038/81497

Weiss T, Miltner W, Adler T, Brückner L, Taub E. Decrease in phantom limb pain associated with prosthesis-induced increased use of an amputation stump in humans. Neurosci Lett. 272: 131–134, 1999. doi: 10.1016/s0304-3940(99)00595-9

Wu C, Kaas J. Reorganization in primary motor cortex of primates with long-standing therapeutic amputations. J Neurosci. 19: 7679–7697, 1999. doi: 10.1523/jneurosci.19-17-07679.1999

